# Prenatal alcohol exposure and facial morphology in a UK cohort

**DOI:** 10.1101/302364

**Authors:** Laurence J Howe, Gemma C Sharp, Gibran Hemani, Luisa Zuccolo, Stephen Richmond, Sarah J Lewis

## Abstract

High levels of prenatal alcohol exposure are known to cause an array of adverse outcomes including foetal alcohol syndrome (FAS); however, the effects of low to moderate exposure are less-well characterised. Previous findings suggest that differences in normal-range facial morphology may be a marker for alcohol exposure and related adverse effects. Therefore, in the Avon Longitudinal Study of Parents and Children, we tested for an association between maternal alcohol consumption and six FAS-related facial phenotypes in their offspring, using both self-report questionnaires and the maternal genotype at rs1229984 in *ADH1B* as measures of maternal alcohol consumption. In both self-reported alcohol consumption (N=4,233) and rs1229984 genotype (N=3,139) analyses, we found no strong statistical evidence for an association between maternal alcohol consumption and facial phenotypes tested. The directions of effect estimates were compatible with the known effects of heavy alcohol exposure, but confidence intervals were largely centred around zero. We conclude that, in a sample representative of the general population, there is no strong evidence for an effect of prenatal alcohol exposure on normal-range variation in facial morphology.

## Introduction

Prenatal alcohol exposure during early embryonic development can have adverse effects on the developing foetus ^1^. The magnitude of the effect is often dependent on the degree of exposure; heavy drinking during pregnancy is associated with the most severe phenotype, foetal alcohol syndrome (FAS) ^2^. Symptoms typically used to diagnose FAS include: stunted anthropometric growth, structural brain anomalies and minor facial anomalies such as a smooth philtrum, a thin upper lip vermillion and shortened palpebral fissures ^3^ In addition to the physically observable symptoms, high levels of prenatal alcohol exposure can cause reduced cognition and psychosocial impairments ^4–6^

While the effects of high levels of prenatal alcohol exposure are well characterised, the effects of lower levels of alcohol exposure are less clear. Individuals exposed to low to moderate levels of alcohol do not typically present with facial, anthropometric or cognitive abnormalities. Indeed, a large systematic review found weak evidence of adverse effects from low to moderate alcohol exposure ^7^ However, previous research has suggested that low to moderate alcohol exposure may have modest adverse effects on offspring cognition and school performance ^8; 9^ and sub-clinical facial differences have been observed between low to moderately exposed, and unexposed individuals ^10^. Variation (within the normal range) in facial morphology may be a marker of alcohol-related cognitive impairments; children exposed to alcohol with more FAS like facial symptoms have been shown to perform worse on psychometric tests ^11^. An association between low levels of alcohol exposure and differential facial morphology would support the hypothesis that low levels of alcohol have adverse effects.

Epidemiological studies investigating the effects of self-reported alcohol intake on health outcomes are problematic because alcohol consumption is often correlated with potential confounders such as tobacco smoking, age and socio-economic status. Indeed, observational studies have reported U- or J-shaped curves where moderate alcohol intake can be associated with improved mortality or morbidity ^12–14^ but follow-up investigations have suggested that these results may be attributable to residual confounding ^13; 15; 16^

Mendelian randomization (MR) is an instrumental variable approach using genetic variants associated with an exposure to explore the relationship between that exposure and an outcome. Assuming a random mode of inheritance, MR may be less susceptible to confounding and reverse causation than observational studies ^17; 18^ The Alcohol Dehydrogenase (ADH) and Aldehyde Dehydrogenase (*ALDH*) genes are two families of genes known to be involved in the production of enzymes that oxidise alcohol ^19^ rs1229984 in *ADH1B* is involved in the metabolism of alcohol to acetaldehyde, individuals with one or more risk alleles are more likely to find drinking unpleasant ^20^ Increased sensitivity to alcohol intake related to this genetic variant has been shown to lead to modified alcohol intake, including alcohol intake in pregnancy ^21^. Although rs1229984 is relatively common in Asian populations ^22^, it is relatively rare in European populations ^21^. Despite this, previous MR studies have successfully utilised this variant as a proxy for alcohol intake in European populations ^8; 16; 23^

In this study, we first investigated whether self-reported low to moderate maternal alcohol intake is associated with facial morphology in the Avon Longitudinal Study of Parents and Children (ALSPAC). Secondly, we used genetic variation in *ADH1B* in an MR framework to estimate the effect of low to moderate maternal alcohol exposure on normal-range facial morphology.

## Methods

### Study participants

#### ALSPAC

We used data on children from the Avon Longitudinal Study of Parents and Children (ALSPAC), a longitudinal study that recruited pregnant women living in the former county of Avon (UK) with expected delivery dates between 1 April 1991 and 31 December 1992. The initial number of enrolled pregnancies was 14,541, which resulted in 14,062 live births and 13,988 children alive at the age of 1. When the oldest children were approximately 7 years of age, the initial sample was boosted with eligible cases who had failed to join the study originally. For analyses of children after the age of 7, the total possible sample size is 15,247 pregnancies, resulting in 14,775 live births. Full details of enrolment have been documented elsewhere ^24–26^ Data was collected from mothers and their partners (during pregnancy and post birth) and from the children (post birth), by self-report questionnaires and clinical sessions. Ethics approval for the study was obtained from the ALSPAC Ethics and Law Committee and the Local Research Ethics Committee. The study website contains details of all available data through a searchable data dictionary (http://www.bristol.ac.uk/alspac/researchers/dataaccess/datadictionary/).

### Measures

#### Facial phenotypes of ALSPAC children

A subset of ALSPAC, consisting of 5,253 children, attended a clinic at the age of 15 years, where high-resolution facial images were taken by Konica Minolta Vivid 900 laser scanners. 4,747 individuals had usable images (506 individuals did not complete the assessment, or the scans were of poor quality and consequently excluded). The derivation of the facial phenotypes are described in more detail in a previous publication ^27^.

The coordinates of 22 facial landmarks were derived using the scans. In this study, facial phenotypes were defined as 3D Euclidean distances between derived facial landmarks. To alleviate multiple testing issues, this study tested 6 facial phenotypes known to be related to the FAS spectrum (**Figure 1**).

**Figure 1:**
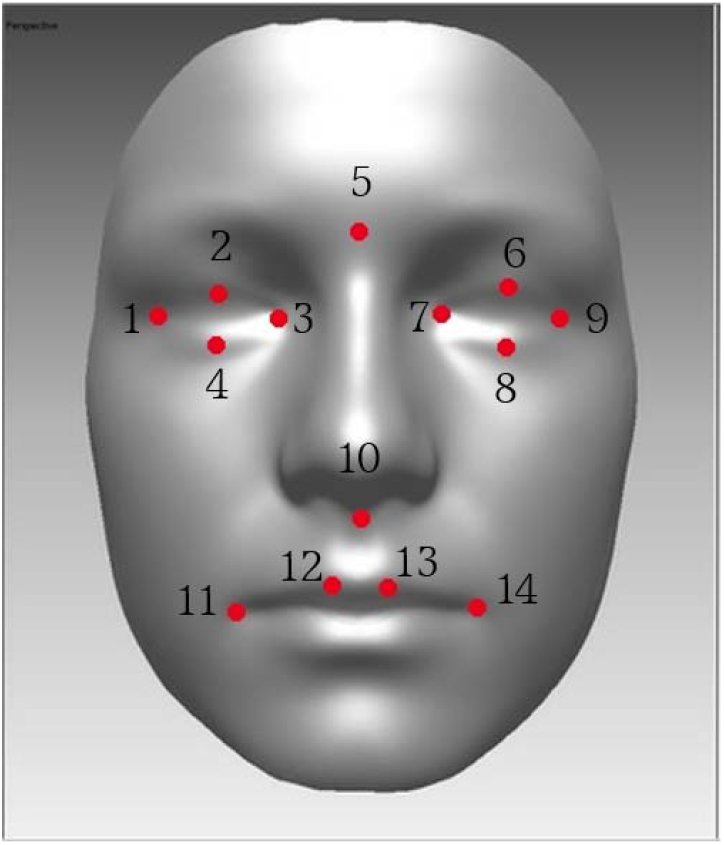
Facial landmarks and phenotypes of interest. This figure shows the 14 facial landmarks used to generate the 6 facial phenotypes. Facial phenotypes were defined as the 3D Euclidean distance between the following landmarks (Average eye palpebral length: 1-3, 7-9; Average eye palpebral width: 2-4, 6-8; Inter-orbital width: 3-7; Nasal length: 5-10; Lip width: 11-14; Philtrum width: 12-13).

#### ALSPAC mothers self-reported alcohol consumption

At around 18 and 32 weeks gestation, ALSPAC mothers completed questionnaires on the average amount and frequency of their alcohol consumption; during pregnancy with alcohol consumption measured in units (roughly 8g of ethanol). Relevant variables, including alcoholic units per week and number of binges in the past month, were used to classify mothers into three categories, based on their self-reported alcohol consumption during pregnancy (non-drinkers during pregnancy, ≤6 units a week and >6 units a week). Further details on questionnaire variables and derivation of classifications are contained in **Supplementary Table 1**.

#### Measurement of potential confounders

Maternal and child data pertaining to covariates included in analyses were measured within the ALSPAC study from questionnaires and clinic sessions.

Maternal age at delivery was calculated using the difference between the mother and child’s dates of birth and gestational age was recorded at birth. Information on maternal smoking was extracted from a questionnaire completed by the mother at 18 weeks gestation; mothers who reported any form of smoking in the first 3 months of pregnancy or in the preceding 2 weeks to the questionnaire were classified as smokers while the non-smokers category included smokers who gave up for pregnancy. Information on maternal education was extracted from a questionnaire completed by the mother at 32 weeks gestation; mothers were asked for their education qualifications and the highest qualification was derived. The height and age of children with available facial scans was recorded at the clinic session.

#### ALSPAC mother’s ADH1B genotype

The *ADH1B* polymorphism rs1229984 was genotyped by KBioscience using the KASPar chemistry (http://www.kbioscience.co.uk/genotyping/genotyping-chemistry.htm). Blind duplicates, plate-identifying repeat samples and Hardy–Weinberg equilibrium tests were used as quality control checks ^21^.

### Statistical analysis

#### Observational analysis

Firstly, to test the association between self-reported maternal alcohol consumption and child facial morphology, we dichotomised mothers into nondrinkers (0 units a week) and mothers reporting alcohol intake (1-6 units a week or >6 units a week). We ran a linear regression of the facial morphology variables on drinking status; adjusting for sex, maternal age, maternal education, maternal smoking and the height and age of the child at the face-shape measurement clinic.

Next, we stratified mothers reporting alcohol intake by alcohol consumption (1-6 units a week and >6 units a week). We re-ran the same analysis, testing differences between the non-drinkers and the two strata separately.

#### Mendelian randomization analysis

For purposes of this analysis, we used the maternal genotype at the SNP rs1229984 in *ADH1B* as a proxy for self-reported maternal intake. Due to the rarity of the homozygous rare genotype, we assumed a dominant effect of the rare allele. To test assumptions about pleiotropy, we used GeneATLAS ^28^ to determine if the SNP was associated with other traits. We then ran a linear regression of the child’s facial morphology variables on the maternal *ADH1B* SNP, adjusting for child’s sex, the first 10 genetic principal components of the mothers, and the height and age of the child at the face-shape measurement clinic.

## Results

### Study sample demographics

#### Observational analysis

Facial phenotype data were available for 4,747 ALSPAC children. We then restricted the sample to maternal-child pairs with complete phenotype data, including; maternal alcohol behaviour during pregnancy, gestational age, maternal age, maternal education, maternal smoking during pregnancy and information on the height and age of the child at the time of the facial scans. The final sample consisted of 4,233 children-mother pairs. More information on the demographics of this sample are contained in **Table 1**.

**Table 1:** Demographics of Observational and Mendelian randomization samples

|  |  | **Observational data-set**<br>**(N = 4233 mother-child pairs)** | **MR data-set**<br>**(N>3000 mother-child pairs)** |
| --- | --- | --- | --- |
| Maternal alcohol behaviour during pregnancy: N (%) | Non-drinkers<br><1-6 units a week<br>>6 units a week | 1302 (30.7)<br>2344 (55.4)<br>587 (13.9) | 927 (30.0)<br>1710 (55.3)<br>454 (14.7) |
| Maternal Education: N(%) | CSE<br>Vocational<br>O level<br>A level<br>University degree | 449 (10.6)<br>329 (7.8)<br>1468 (34.7)<br>1224 (28.9)<br>763 (18.0) | 298 (9.6)<br>227 (7.3)<br>1027 (33.2)<br>945 (30.5)<br>591 (19.1) |
| Maternal smoking during pregnancy: N (%) | Yes<br>No | 694 (16.4)<br>3539 (83.6) | 508 (16.2)<br>2631 (83.8) |
| Gestational age: (weeks) | Mean (SD) | 39.5 (1.8) | 39.5 (1.8) |
| Estimated age of mother at conception: (years) | Mean (SD) | 28.5 (4.5) | 28.6 (4.4) |
| Child's sex: N (%) | Male<br>Female | 2003 (47.3)<br>2230 (52.7) | 1498 (47.2)<br>1678 (52.8) |
| Age of child at clinic: (years) | Mean (SD) | 15.4 (0.3) | 15.4 (0.3) |
| Height of child at clinic: (cm) | Mean (SD) | 169.2 (8.4) | 169.3 (8.4) |
| Average eye palpebral width: (mm) | Mean (SD) | 11.6 (1.4) | 11.6 (1.4) |
| Average eye palpebral length: (mm) | Mean (SD) | 27.3 (1.8) | 27.2 (1.8) |
| Lip width: (mm) | Mean (SD) | 53.1 (4.0) | 53.1 (4.0) |
| Philtrum width: (mm) | Mean (SD) | 12.9 (1.9) | 12.8 (1.9) |
| Nasal length: (mm) | Mean (SD) | 52.4 (2.9) | 52.5 (2.9) |

#### Mendelian randomization analysis

Again, starting with the 4,747 ALSPAC children with facial phenotype data, we selected maternal-child pairs with the maternal *ADH1B* SNP genotyped and information on the height and age of the child at the time of the facial scans. The final sample consisted of 3,139 child-mother pairs. More information on this sample is contained in **Table 1**.

### Observational analysis

We did not find strong statistical evidence for an association between selfreported maternal alcohol consumption and the 6 facial phenotypes tested, although the directions of effect in our results were compatible with the symptoms of FAS for facial phenotypes tested (**Table 2**). In the stratified analysis, effect sizes of higher magnitude were observed in the >6 units a week for 5 out of 6 phenotypes, suggestive of a possible dose-response relationship. However, in the stratified analysis we also did not find strong evidence of an association between alcohol exposure and facial morphology (**Supplementary Table 2**).

**Table 2:** Association of maternal self-reported alcohol intake and maternal *ADH1B* genotype with child’s facial phenotypes

| Facial morphology variable | Non-drinkers against any alcohol intake (<1-6 units a week or >6 units a week) (N=4233) <sup>1</sup> |  |  | *ADH1B* Mendelian randomization analysis Adjusted for 10 PCs, height, age, sex (N=3139) <sup>2</sup> |  |  |
| --- | --- | --- | --- | --- | --- | --- |
|  | Beta (95% C.I.) | P-value | Direction concordant with literature | Beta (95% C.I.) | P-value | Direction concordant with literature |
| Average eye palpebral width ps_pi | -0.03 (-0.12, 0.06) | 0.51 | Yes | -0.07 (-0.30, 0.15) | 0.52 | Yes |
| Average eye palpebral length ex_en | -0.06 (-0.17, 0.06) | 0.31 | Yes | -0.20 (-0.50, 0.10) | 0.19 | Yes |
| Lip width chL_chR | 0.06 (-0.20, 0.32) | 0.67 | Yes | 0.17 (-0.52, 0.85) | 0.63 | Yes |
| Philtrum width cphL_cphR | -0.03 (-0.15, 0.09) | 0.65 | Yes | -0.16 (-0.47, 0.16) | 0.33 | Yes |
| Nasal length sn_n | -0.04 (-0.23, 0.16) | 0.71 | Yes | -0.10 (-0.60, 0.39) | 0.68 | Yes |
| Inter-orbital width enL_enR | 0.17 (-0.01, 0.34) | 0.058 | Yes | 0.06 (-0.38, 0.51) | 0.87 | Yes |

<sup>1</sup> Covariates: Sex, Gestational age, Maternal age, Maternal education, Maternal smoking, Child's height and Child's age at Face-shape measurement clinic
<sup>2</sup> Covariates: Sex, Principal components, Child's height and Child's age at Face-shape measurement clinic

### Mendelian randomization analysis

#### Maternal ADH1B SNP, alcohol behaviour and pleiotropy

We confirmed that the *ADH1B* SNP was strongly predictive of alcohol behaviour in our sample; one or more of the rarer A alleles of rs1229984 was associated with reduced odds of reported maternal drinking relative to no drinking OR: 0.54 (95% C.I. 0.38, 0.75; P < 0.001). While self-reported maternal drinking was associated with all four potential confounders (maternal education, smoking, gestational age at delivery and maternal age), there was weak evidence for an association between these variables and the *ADH1B* SNP (**Supplementary Table 3**).

In the GeneATLAS, based on the UK Biobank, the *ADH1B* SNP was strongly associated with alcohol intake frequency (P = 1.59 × 10^−148^). However, in this much larger sample, there was some evidence that the SNP is pleiotropic. Notably there is strong evidence that the SNP is associated with socio-economic status markers such as the Townsend deprivation index (P = 1.98 × 10^−7^) and number of vehicles in household (4.8 × 10^−9^), as well as measures of adiposity such as body mass index (P = 3.65 × 10^−11^).

#### Maternal ADH1B genotype and child’s facial morphology

We then tested for association between the maternal *ADH1B* SNP and 6 facial phenotypes. We found no strong statistical evidence that the maternal *ADH1B* SNP was associated with the 6 facial phenotypes, but we did find that the directions of effect were consistent with the symptoms of FAS, as in the observational analysis (**Table 2**).

## Discussion

In this study, we used self-reported maternal alcohol intake and the maternal genotype for rs1229984 in *ADH1B* to test the hypothesis that alcohol exposure has a detectable effect on normal-range facial variation in the general population. In both observational and MR analyses, we found weak statistical evidence for an association. The directions of effect were compatible with the known effects of heavy alcohol exposure, but confidence intervals were wide. The implication of these results is that there is no strong evidence for an effect of alcohol exposure on the FAS-related facial phenotypes tested in our study sample.

Our findings are consistent with the well-characterised difficulties of identifying the effects of prenatal alcohol exposure ^11^ as well as with the findings of a previous systematic review ^7^. Contrastingly, a previous investigation by Muggli and colleagues found evidence of sub-clinical facial differences at 12 months between offspring of non-drinking mothers and offspring of light to moderate drinking mothers ^10^ The discordance in results may be explained by differences in the timing and complexity of facial morphology measurements used in both studies; the study by Muggli and colleagues measured facial morphology at age 12 months using a 3-D surface registration algorithm consisting of 69,587 points, whereas our study measured facial morphology using six Euclidean distances at age 15. Poor statistical power is unlikely to explain the lack of replication, because the sample size in our study was more than 10 times that of the study by Muggli and colleagues.

A considerable strength of this study is the use of triangulation of methods ^29^; using both self-reported alcohol intake and the *ADH1B* SNP as measures of alcohol exposure and finding concordant results between the two methods. A further advantage is the substantially large sample size used in our analyses, compared to similar previous studies ^10; 30–32^. However, although our MR analysis included over 3000 mother-child pairs, it may still lack statistical power because of the low frequency of the rs1229984 minor allele. A further limitation of the study is that both the observational and MR analyses may be susceptible to confounding. In theory, MR analyses are less affected by confounding but there is some evidence that the *ADH1B* SNP may be associated with potential confounders such as social class and education ^16^ However, in this instance, any confounding related to socio-economic factors would likely bias results away from the null hypothesis and so do not affect our conclusions. Our use of relatively simple facial phenotypes compared to a previous study ^10^, that constructed phenotypes from thousands of facial landmarks, may also be a limitation. More detailed facial phenotyping may better capture surface topography and overall facial shape, although it is worth noting that some of the phenotypes tested in this study are included in the latest diagnostic criteria for FAS ^33^ A further limitation is that when estimating the effect of maternal genotypes using MR, the child’s genotypes may be a confounder if they also affect the phenotype of interest ^34^

To conclude, in a large sample size of children, we found no strong evidence for an association between maternal alcohol consumption and facial morphology of their offspring. A lack of statistical power hinders stronger conclusions, future metaanalyses could allow more definitive conclusions to be made. The absence of strong evidence for an effect does not necessarily imply that there is no effect, but given our well-powered observational analysis, it suggests that the effect size is likely to be small. Our results have important implications for the potential use of facial morphology to identify individuals adversely affected by prenatal alcohol exposure. Further work in ALSPAC could involve the use of more complex facial phenotypes, that may better identify facial-structure differences ^10; 35^

## Acknowledgements

We are extremely grateful to all the families who took part in this study, the midwives for their help in recruiting them, and the whole ALSPAC team, which includes interviewers, computer and laboratory technicians, clerical workers, research scientists, volunteers, managers, receptionists and nurses.

